# *ascend*: R package for analysis of single cell RNA-seq data

**DOI:** 10.1101/207704

**Authors:** Anne Senabouth, Samuel W Lukowski, Jose Alquicira Hernandez, Stacey Andersen, Xin Mei, Quan H Nguyen, Joseph E Powell

## Abstract

**Summary:** *ascend* is an R package comprised of fast, streamlined analysis functions optimized to address the statistical challenges of single cell RNA-seq. The package incorporates novel and established methods to provide a flexible framework to perform filtering, quality control, normalization, dimension reduction, clustering, differential expression and a wide-range of plotting. ascend is designed to work with scRNA-seq data generated by any high-throughput platform, and includes functions to convert data objects between software packages.

**Availability:** The R package and associated vignettes are freely available at https://github.com/IMB-Computational-Genomics-Lab/ascend.

**Supplementary information:** An example dataset is available at ArrayExpress, accession number E-MTAB-6108

## 1 Introduction

Single cell RNA sequencing (scRNA-seq) has revolutionised the way we understand differences in transcriptional programs. Recent advances in droplet-based molecular biology techniques, coupled with microfluidics have given us the ability to examine the transcriptomes of tens of thousands of single cells simultaneously. This has enabled the characterization of gene expression variation for each cell in a heterogeneous population. Platforms and protocols such as 10X Genomics Chromium [1], Drop-seq [2], InDrop [3], and others (reviewed in [4]) provide accessible high-throughput methods for generating scRNA-seq data. This breakthrough in technology has generated new challenges in data management, normalization, statistical methods, data visualization, and computing limitations because current methods for standard RNA-seq analysis cannot be readily applied to scRNA-seq data.

Here we present *ascend*, an R package designed to create a simple and streamlined workflow for the analysis of scRNA-seq experiments. *ascend* is designed to handle data generated from any single cell library preparation platform; which can include data from either single and pair-end reads, and including or not including unique molecular identifiers (UMIs). Software such as *scpipe* (https://github.com/LuyiTian/scPipe) [5] provides a unified bioinformatics pipeline to generate a count matrix from raw sequence data, but does not include any further functionality for down-stream analysis of scRNA-Seq data. *ascend* imports scRNA-seq data following bioinformatics processing, performs user-friendly quality control, filtering, normalisation, dimension reduction, clustering, differential expression and visualization. It includes functions to leverage multiple CPUs, allowing most analyses to be performed on a standard desktop or laptop.

## 2 *ascend*

### 2.1 Data object

The core of the *ascend* R package is the Expression and Metadata Set (EMSet), an S4 object made specifically for scRNA-seq experiments requiring only a user-supplied gene-cell expression matrix. Metadata such as cell- or gene-related information, including custom controls and spike-ins, can be incorporated. We have included a specific function to convert a GeneBCMatrix object generated by the 10X Genomics Cell Ranger pipeline into an EMSet object. Class-specific convenience functions allow object components to be manipulated in place or extracted with ease and standard R functions can be used to modify metadata. In addition, to promote cross-package compatibility, a convenience method is available to convert the EMSet to another common class, the SCESet, created by the R packages *scater* [6] and *scran* [7].

### 2.2 Filtering and quality control

Cell-specific and gene-specific metrics are generated upon the creation of an EMSet and are recalculated when changes to the object are made. ascends filtering functions identify and remove low-quality cells based on library size and gene expression. Filtering of outliers greater than a specified mean absolute deviation (MADs) is based on the workflow described by Lun *et al.* [7], and cells expressing a high percentage of defined control genes (e.g. mitochondrial/ribosomal) can also be excluded. Customised sets of control genes, such as spike-ins, can be substituted. Finally, genes detected in a low percentage of cells can be excluded. Functions include default parameters, which can be edited by a user.

### 2.3 Removal of batch effects and normalisation

Analysis of large-scale scRNA-seq data often requires combining multiple samples. Therefore, to remove systematic biases due to technical variation between batches, ascend removes potential batch effects at two levels, between cells and between samples. Here, the effects of confounding factors, such as experimental conditions or cell cycle genes can also be removed. Cell-cell normalization is another crucial step to remove technical variation between individual cells. ascend offers two options to calculate a normalization size factor for each cell: a computationally intensive deconvolution method [8], and we adjusted the RLE approach (Relative Log Expression, as described in Anders and Huber, 2010) to find geometric means in zero-inflated data (refer to *ascend* vignettes for more details). The former method takes into account the zero-inflation issue by pooling cells, while the latter uses gene expression values higher than zero to calculate the geometric mean of a gene. After normalization, the data is ready for analysis to find biological changes/differences.

### 2.4 Reduction of high-dimensional space

Dimensionality reduction is an important step for scRNA data analysis, since scRNA data is multiple orders of magnitude larger than bulk RNA-seq data (*n*-cells * *m*-genes). Moreover, the expression levels of many genes are likely to be correlated, therefore the problem of (multi)collinearity is common, while and additional factors such as dropouts, and high expression variation increase the noise in the data. *ascend* implements principal component analysis (PCA) or multidimensional scaling (MDS) to reduce the dimensions of the data and preserve the data structure (i.e. explain most of the variance between cells). A *t*-SNE (*t*-distributed Stochastic Neighbour Embedding) procedure is used only to visualise cells in a low-dimensional space labeled by batch, cluster or the expression levels of specific genes.

### 2.5 Clustering

Clustering cells into subpopulations is a crucial step that determines downstream analysis in singlecell workflow [9,10]. *ascend* implements our CORE algorithm (Nguyen *et al.*, 2017, under review) as an unsupervised approach that does not require user defined-parameters, and a stable, optimal number of clusters is computed automatically. The method is fast and scalable, enabling the detection of small clusters at high resolution as rare subpopulations, or larger clusters for more general classification with simpler downstream analysis.

Briefly, from the normalised gene expression, a Euclidean distance matrix between cells is calculated. An unsupervised dendrogram is then constructed by applying hierarchical clustering as described (Nguyen *et al.*, under review). The dendrogram is dynamically reclustered by a top-down split and merging process over multiple iterations with changing tree-height thresholds. This approach merges smaller clusters into larger consensus clusters, and uses an adjusted Rand index to compare different clustering results to find the optimal number of clusters.

### 2.6 Differential expression

After decomposing the dataset into subpopulations, *ascend* provides functionalities to compare these subpopulations for finding biological signatures that distinguish them. The analysis is based on negative binomial tests from DESeq [11]. We introduced several modifications that allow (i) more accurate estimation of fold change (adjusted fold change), and (ii) more efficient multiprocessing (divide and conquer approach) to handle large datasets.

## 3 Conclusion

In summary, *ascend* is a user-friendly and computationally efficient package for analysing scRNA-seq data from all experimental platforms. *ascend* implements a state-of-the-art unsupervised clustering method and integrates established analysis techniques for normalisation and differential gene expression. The *ascend* package and context-specific tutorials addressing a range of analytical scenarios are available at https://github.com/IMB-Computational-Genomics-Lab/ascend.

**Figure 1:**
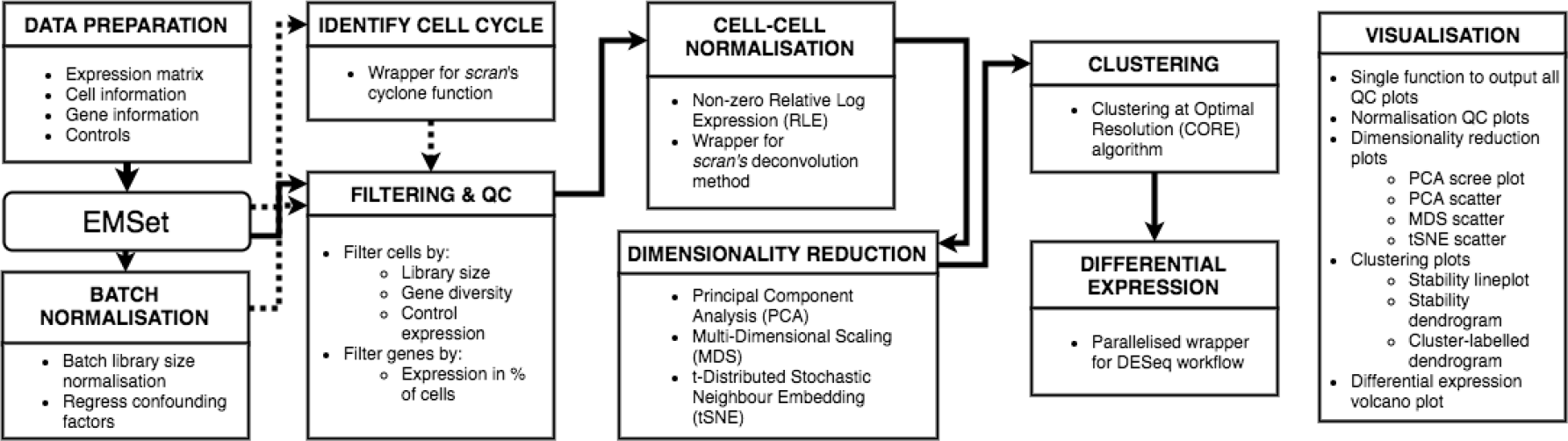
The *ascend* workflow. Solid lines indicate general workflow and dashed lines indicate optional steps. Visualisation can be performed after the dimensionality reduction or clustering steps.

## Acknowledgements

The authors wish to thank the developers of open source single cell research software, in particular Davis McCarthy, Aaron Lun, and Luke Zappia.

